# MSDC-0602K, A Novel Insulin-Sensitizer Improves Insulinemia and Fatty Liver Disease Alone and in Addition to Liraglutide in Mice

**DOI:** 10.1101/2020.12.10.420190

**Authors:** Dakota R. Kamm, Kelly D. Pyles, Martin C. Sharpe, Laura Healy, Jerry R. Colca, Kyle S. McCommis

## Abstract

**Aims/hypothesis:** Insulin sensitizers and incretin mimetics are antidiabetic agents with vastly different mechanisms of action. Additionally, while thiazolidinedione (TZD) insulin sensitizers are associated with the side-effect of weight gain, glucagon-like peptide-1 receptor agonists (GLP-1RAs) can induce weight loss. We hypothesized that combination therapy with a novel TZD insulin sensitizer and the GLP-1RA Liraglutide would more significantly improve mouse models of diabetes and nonalcoholic steatohepatitis (NASH) compared to individual therapy.

**Methods:** *db/db* mice were treated with the novel TZD MSDC-0602K by oral gavage, Liraglutide (Lira) by s.c. injection, combination 0602K+Lira, or both vehicle solutions. Vehicle-treated *db/*+ mice were included as non-obese controls. To assess treatment effects on nonalcoholic fatty liver disease, MS-NASH mice were similarly treated with vehicle, either drug individually, or in combination. Lastly, islets were isolated from C57BL/6J mice to assess glucose-stimulated insulin secretion (GSIS).

**Results:** 0602K-treated *db/db* mice displayed slight weight gain but completely corrected glycemia and markedly improved glucose tolerance. Lira slightly reduced body weights and modestly improved glycemia. 0602K+Lira combination still induced slight weight gain but completely corrected glycemia and improved glucose tolerance beyond lean *db/*+ levels. As expected, 0602K resulted in reduced plasma insulin, whereas Lira further increased the hyperinsulinemia of *db/db* mice. Surprisingly, 0602K+Lira treatment reduced plasma insulin and C-peptide to the same extent as mice treated with 0602K alone. 0602K did not directly reduce GSIS in isolated islets, thus the reduced insulinemia with 0602K is likely compensatory due to improved insulin action. In the MS-NASH mouse model, both 0602K or Lira alone improved plasma ALT and AST, and liver histology, but more significant improvements were observed with 0602K+Lira combination therapy. 0602K or 0602K+Lira also increased pancreatic insulin content in both *db/db* and MS-NASH mice.

**Conclusions:** MSDC-0602K corrected glycemia and reduced insulinemia when given alone, or in combination with Lira. However, 0602K+Lira combination more significantly improved glucose tolerance in *db/db* mice, and more significantly improved liver histology in MS-NASH mice.

## Introduction

Insulin resistance can result in elevated rates of insulin secretion and hyperinsulinemia. If unresolved, pancreatic beta cell failure can lead to loss of beta cell mass via cell death or dedifferentiation [1, 2]. Insulin sensitizers are an attractive therapeutic strategy as they not only target the core defective insulin signaling pathways but can also reduce this stress on beta cells and preserve beta cell mass [3]. However, clinical use of the main class of insulin sensitizers, the thiazolidinediones (TZDs), is limited due to side effects such as weight gain, edema, bone loss, and bladder cancer risk. These side effects are thought to be due to agonism of the nuclear receptor peroxisome proliferator-activated receptor γ (PPARγ). Yet numerous studies have described acute, PPARγ-independent effects of TZDs [4-7], which have led to the development of several TZDs with very low affinity for PPARγ, such as MSDC-0602K and MSDC-0160. The molecular target of these compounds is the mitochondrial pyruvate carrier (MPC) [8-10], which is also inhibited by the traditional PPARγ-activating TZDs [9]. The clinical profile of PPARγ-sparing TZDs appears improved compared to traditional TZDs with respect to edema, bone loss, and degree of weight gain [11-13].

Another popular class of antidiabetic agents is the glucagon-like peptide-1 receptor agonists (GLP-1RA). These incretin-like peptides improve glycemia by increasing postprandial insulin secretion, suppressing glucagon secretion, and delaying gastric emptying. GLP-1RAs such as Liraglutide can also induce weight loss in humans and animal models [14, 15].

Insulin resistance and diabetes are driving factors for the development and progression of nonalcoholic fatty liver disease (NAFLD) and nonalcoholic steatohepatitis (NASH). While there are currently no approved therapies for treatment of NAFLD/NASH, both TZDs [13, 16-21] and GLP-1RAs [22-25] can improve aspects of liver pathology in humans and animal models. The purpose of this current study was to investigate whether combining the PPARγ-sparing TZD MSDC-0602K and the GLP-1RA Liraglutide would better improve mouse models of diabetes and NASH.

## Methods

All experiments and procedures were approved by the Institutional Animal Care and Use Committee of Saint Louis University, and conform to NIH guidelines for the care and use of laboratory animals [26]. All mice were housed in standard rodent caging at 20-25°C with *ad libitum* access to food and water with a light cycle from 6:00AM – 6:00PM.

### *db/db* mouse studies

Five-week-old male *db/db* mice on C57BL/6J background, and age/sex matched *db/*+ control mice were purchased from The Jackson Laboratory (B6.BKS(D)-*Lepr*^db^/J, stock #000697; Bar Harbor, ME, USA). Mice were provided *ad libitum* access to water and chow (Laboratory Rodent Diet 20 (5L0B), LabDiet, St. Louis, MO, USA). At 9 weeks of age, blood glucose was measured by nick of the tail vein and measuring with a Contour Next glucometer (Ascensia Diabetes Care, Parsippany, NJ, USA). *db/db* mice were then divided into treatment groups based on averaged body weights and blood glucose concentrations: Vehicle (Veh), 30mg/kg MSDC-0602K (0602K) gavage daily, 0.2mg/kg Liraglutide (Lira) s.c. injection every-other-day, or combined 0602K+Lira. Lira was obtained from MedChemExpress (HY-P0014; Monmouth Junction, NJ, USA). Vehicle gavage solution was 1% low-viscosity carboxymethylcelluse, 0.1% Tween-80, and 1% DMSO. 0.9% NaCl solution was vehicle for s.c. injection. *db/*+ and *db/db* Veh mice received both the gavage and s.c. injection vehicles. 0602K-treated mice also received vehicle s.c. injection while Lira-treated mice also received gavage vehicle. After 3 weeks of treatment, mice were euthanized by CO_2_ asphyxiation and blood collected by cannulation of the inferior vena cava, placed into an EDTA-containing tube, and centrifuged at 8,000 x g for 8 minutes at 4°C to collect plasma. Tissues were dissected, blotted dry, frozen in liquid nitrogen, and stored at -80°C. Pieces of liver and pancreas were placed in neutral buffered formalin for histological evaluation.

### Glucose tolerance test

After 1.5 weeks of treatment, mice were challenged by a GTT. At 8:00AM, mice were weighed and fasted. 4 hours later, mice were injected i.p. with 1g/kg D-glucose in 0.9% NaCl immediately after measuring blood glucose (T=0) by a nick to the tail vein and a Contour Next glucometer (Ascensia Diabetes Care, Parsippany, NJ, USA). Blood glucose was monitored at 15, 30, 60, 90, and 120 minutes post-injection by removing the scab from the tail.

### MS-NASH mouse studies

Studies of the MS-NASH mouse model (Jackson Laboratory MSNASH/PcoJ, stock #030888) were performed at CrownBiosciences Inc. Beginning at 9 weeks of age, male MS-NASH mice were fed *ad libitum* with a “western diet” (D12079B, Research Diets Inc., New Brunswick, NJ, USA) containing 40% kcal fat, 17% kcal protein, 43% kcal carbohydrate, and 1.5g/kg cholesterol, and *ad libitum* drinking water containing 5% (w/v) fructose. Body weight was measured weekly and blood was collected via the tail vein for measurement of blood glucose and plasma alanine aminotransferase (ALT) and aspartate aminotransferase (AST) every 4 weeks as described below. Based on average ALT concentrations, mice were divided into 4 treatments: Veh, 0602K, Lira, or 0602K+Lira, performed identically to the *db/db* studies described above for a duration of 6 weeks. Mice were euthanized by CO_2_ asphyxiation and blood collected via cardiac puncture. Tissues were dissected and frozen in liquid nitrogen. Pieces of liver and pancreas were fixed in neutral buffered formalin for histological examination.

### RIPCreMPC2-/-mouse studies

Generation of *Mpc2* floxed mice and RIPCreMPC2-/-pancreatic beta cell MPC2-/-mice on the C57BL/6J background were reported previously [10, 27]. For these studies, 6-week-old male RIPCreMPC2-/- and littermate fl/fl control mice were fed 60% high-fat diet for 10 weeks (D12492, Research Diets Inc., New Brunswick, NJ, USA). Based on body weight, mice were divided into two single gavage treatments: Vehicle, or 30mg/kg 0602K. 16h post gavage, mice were euthanized by CO_2_ asphyxiation. Blood was collected from the inferior vena cava into an EDTA-coated tube, processed as above, and plasma stored at -80°C.

### Pancreatic islet isolation and glucose-stimulated insulin secretion assay

Wildtype male and female C57BL/6J mice were obtained (Jackson Laboratory, stock #000664) and maintained on normal chow. Between 10-20 weeks of age, mice were euthanized by 3-5% isoflurane inhalation and cervical dislocation. Islets were isolated similar to previously described procedures [27, 28], after perfusion of the pancreas through the common bile duct with 5-10mL of calcium-free Hanks-buffered saline solution (HBSS) supplemented with 0.4mg/mL Type V collagenase (C9263, Sigma, St. Louis, MO, USA). After overnight culture, islets were treated with either 1 or 23mM glucose and either DMSO vehicle or 10μM 0602K in a 37°C incubator for 1h. Insulin concentration of the supernatants was measured with a mouse/rat insulin ELISA (EZRMI-13K, EMD Millipore, Billerica, MA, USA).

### Plasma hormone, metabolite, and ALT and AST assays

Plasma fructosamine levels were measured with a colorimetric kit (K450-100, BioVision Inc., Milpitas, CA, USA). Plasma insulin and C-peptide were measured with mouse/rat ELISAs (EZRMI-13K and EZRMCP2-21K, respectively, EMD Millipore, Billerica, MA, USA). Plasma non-esterified fatty acids (NEFA) were measured by enzymatic colorimetric assay (NEFA-HR(2), FUJIFILM Wako, Mountain View, CA, USA). Plasma triacylglycerol (TAG) and cholesterol were measured with colorimetric assays (TR22421 and TR13421, respectively, ThermoFisher Scientific, Middletown, VA, USA). Plasma ALT and AST concentrations were measured with kinetic spectrophotometric assays (A524-150 and A559-150, respectively, Teco Diagnostics, Anaheim, CA, USA).

### Liver TAG and glycogen assays

Liver TAG concentrations were measured as performed previously [16]. 40-140mg of frozen tissue was homogenized using a bead homogenizer (Mini-Beadbeater, Biospec Products Inc., Bartlesville, OK, USA) in 0.9% NaCl at a volume to provide 0.1mg liver/μL. Liver homogenate was combined (1:1) with 1% sodium deoxycholate, vortexed, and placed at 37°C for 5min to solubilize lipids. TAG was then measured by colorimetric assay (TR22421, ThermoFisher Scientific, Middletown, VA, USA).

Liver glycogen was measured using previously described methods [29]. 20-75mg of liver tissue was boiled in 300μL of 30% KOH at 100°C for 30min. Tubes were cooled on ice and 100μL of 2% Na_2_SO_4_ and 800μL of 100% EtOH added and tubes vortexed. Tubes were boiled for 5min and centrifuged at 16,000 x g for 5min then supernatant aspirated. The pellet was dissolved in 1mL of 80% EtOH and recentrifuged 16,000 x g for 5min. Final pellets were resuspended in 200μL of 0.3mg/mL amyloglucosidase (Sigma, St. Louis, MO, USA) in 0.2M sodium acetate. Serial dilutions of 10mg/mL oyster glycogen (Sigma, St. Louis, MO, USA) were prepared as standards. Samples and standards were incubated in a 40°C water bath for 3h, then diluted 1:1 with H_2_O, and 5μL of each added to a 96-well plate. 200μL of glucose assay buffer (0.3 M Triethanolamine, pH∼7.5, 4mM MgCl_2_, 2mM ATP, 0.9mM NADP+, and 5μg/mL Glucose-6-phosphate dehydrogenase) was added to each well, and absorbance measured at 340nm. 1μg of Hexokinase (Sigma, St. Louis, MO, USA) was then added to each well, the plate incubated at room temperature in the dark for 30 min, and absorbance remeasured at 340nm.

### Pancreas and liver histology and immunohistochemistry analyses

Formalin fixed liver and pancreas sections were embedded in paraffin blocks and sectioned onto glass slides. Liver slides were stained for H&E and Sirius Red. Digitized liver slides were evaluated by a histopathologist blinded to mouse treatment and group designations, and scored for NAFLD activity (combined scores of steatosis, inflammation, and hepatocyte ballooning) and fibrosis similar to human biopsy scoring [30].

For the *db/db* study, unstained pancreas slides were used for immunofluorescence as performed previously [27]. Slides were rehydrated then permeabilized with 1mg/mL trypsin (T7168, Sigma, St. Louis, MO, USA) in H_2_O for 25min at room temperature. Slides were washed 2 x 5min with 0.2% NP40-PBS, and blocked with 0.2% NP40-PBS containing 3% BSA for 30min at room temperature. Slides were incubated with primary antibodies overnight at 4°C in a humidity chamber. Primary antibodies were guinea pig anti-insulin polyclonal and mouse anti-glucagon monoclonal, both 1:100 (ab7842 and ab10988, respectively, Abcam, Cambridge, MA, USA). The following day, slides were washed 3 x 5min with 0.2% NP40-PBS, then incubated with secondary antibodies for 2h at room temperature. Secondary antibodies were goat anti-guinea pig, Alexa Fluor488 and goat anti-mouse, Alexa Fluor594 (A11073 and A21125, respectively, ThermoFisher Scientific, Middletown, VA, USA). Slides were washed 3 x 5min with 0.2% NP40-PBS, and glass coverslips mounted with ProLong Diamond with DAPI (P36971, ThermoFisher Scientific, Middletown, VA, USA). Slides were imaged on a Leica DM5500B fluorescence microscope with Leica DFC365 FX camera (Leica Microsystems Inc., Buffalo Grove, IL, USA). Intensity of the green insulin fluorescence of pancreatic islets relative to nearby exocrine pancreas was measured using NIH Image J software.

For the MS-NASH study, pancreas slides were stained for H&E or prepared for insulin immunohistochemistry by performing heat-induced epitope retrieval using EnVision FLEX Target Retrieval Solution, low pH∼6 (Dako, K8005; Agilent, Santa Clara, CA, USA), with preheating to 80°C and increased temperature to 95°C for 20min after slides were added. Slides were then incubated in 3% H_2_O_2_ for 5min and incubated with rabbit anti-Insulin antibody, 1:100 for 45min (#4590, Cell Signaling Technology, Danvers, MA, USA). Slides were then washed and secondary antibody was EnVision+ anti-rabbit labelled polymer-HRP applied for 30min (Dako, K4003; Agilent, Santa Clara, CA, USA), then DAB+ chromogen solution for 5min. Slides were then rinsed in water and counterstained with modified Harris hematoxylin (Dako, S3301; Agilent, Santa Clara, CA, USA). Slides were cover-slipped and scanned at 20X on an Aperio Scanscope XT (Leica Biosystems, Buffalo Grove, IL, USA), and insulin+ cells quantified by HALO software (Indica Labs, Albuquerque, NM, USA).

### RNA isolation, cDNA synthesis, and quantitative RT-PCR analyses

5-20mg of frozen liver tissue was homogenized in 1mL of RNA-STAT reagent (Tel-Test, Friendswood, TX, USA) with isopropanol and EtOH precipitation. RNA pellets were resuspended in 200uL water and assessed by Nanodrop (ThermoFisher Scientific, Middletown, VA, USA). 1ug of RNA was reverse transcribed into cDNA by Superscript VILO kit (ThermoFisher Scientific, Middletown, VA, USA), using an Applied Biosystems 2720 Thermal Cycler (ThermoFisher Scientific, Middletown, VA, USA). qPCR was performed for all samples in duplicate with Power SYBR Green using an Applied Biosystems QuantStudio-3 real-time thermocycler (ThermoFisher Scientific, Middletown, VA, USA). Target gene Ct values were normalized to reference gene (*Rplp0*) Ct values by the 2^-ΔΔCt^ method. Oligonucleotide primer sequences are listed in electronic supplementary material Table 1.

### Statistics

All data are expressed as means±SEM, with individual data points shown as dot plots, or curves over time displayed as means±SEM. Individual data points represent a single mouse. No data were excluded from any study measurements. Investigators were not blinded to genotype or treatment. The *db/db* study was performed with 3 separate cohorts/replicates of mice and combined. For liver histology analysis of the MS-NASH studies, the pathologist was blinded to mouse number and treatment groups. The MS-NASH study was performed with a single cohort of mice. Statistical analyses were performed using GraphPad Prism 8 (GraphPad, San Diego, CA, USA) using ordinary one-way ANOVA and Tukey’s multiple comparisons tests and *p*<0.05 considered statistically significant.

## Results

### Effects of MSDC-0602K and Liraglutide on body weight, tissue weights, and glycemia of *db/db* mice

Beginning at 9 weeks of age, *db/db* mice were treated with either MSDC-0602K, Liraglutide, combination, or both vehicle solutions. 0602K-treatment caused slight weight gain and Lira-treatment caused slight weight loss compared to vehicle-treated *db/db* mice (Fig. 1a). Surprisingly, mice treated with 0602K+Lira combination displayed slight weight gain similar to mice treated with 0602K alone (Fig. 1a). Blood glucose was monitored weekly and were completely corrected to *db/*+ levels by 0602K or 0602K+Lira treatments, whereas Lira treatment more modestly improved glycemia (Fig. 1b). Both drugs individually improved glucose tolerance compared to vehicle-treated *db/db* mice, and 0602K+Lira combination improved glucose tolerance beyond even lean *db/*+ mouse levels (Fig. 1c,d). At sacrifice, 4h fasted blood glucose levels were completely corrected by 0602K or 0602K+Lira (Fig. 1e). Plasma fructosamine levels supported the weekly glucose measurements indicating that Lira modestly improved glycemia while 0602K or 0602K+Lira completely corrected glycemia (Fig. 1f). These results indicate that while 0602K alone is able to correct glycemia, the combination of 0602K+Lira better improves glucose tolerance in *db/db* mice.

**Fig. 1.**
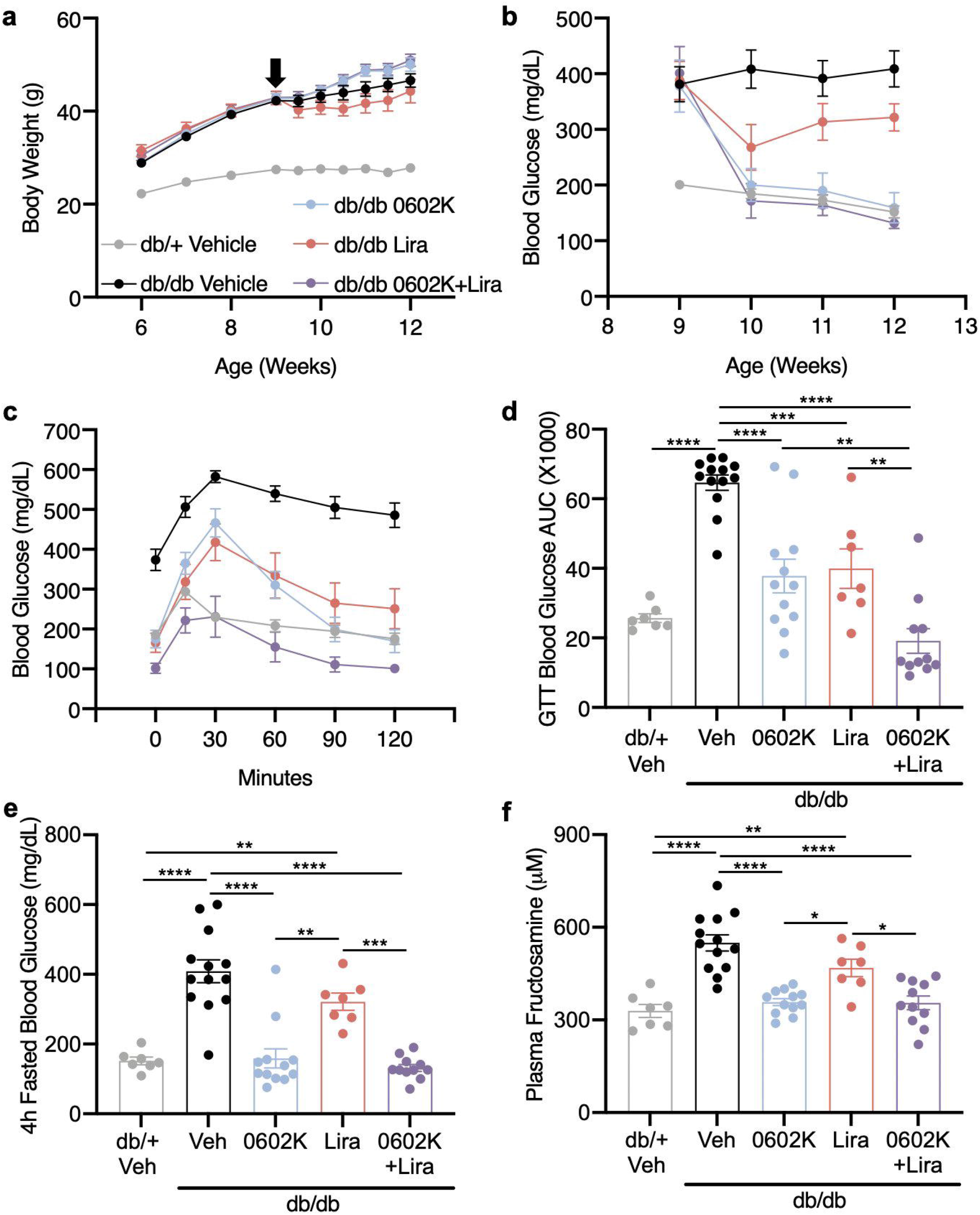
MSDC-0602K alone and in combination with Liraglutide improves glycemia and glucose tolerance in *db/db* mice. (**a**) Average weekly body weights of each treatment group. (**b**) Average weekly blood glucose concentrations. (**c**,**d**) Blood glucose excursions and calculated area under the curve from an i.p. glucose tolerance test (GTT). (**e**) 4h fasted blood glucose concentrations measured at time of sacrifice. (**f**) Plasma fructosamine concentrations. n= 7 *db/*+ Veh; 13 *db/db* Veh, 12 *db/db* 0602K, 7 *db/db* Lira, and 11 *db/db* 0602K+Lira. All data are expressed as means±SEM or means±SEM within dot plot of individual data points. Individual data points represent a single mouse. Ordinary one-way ANOVA with Tukey’s multiple comparison’s test: \**p*<0.05, \*\**p*<0.01, \*\*\**p*<0.001, and \*\*\*\**p*<0.0001.

Epididymal white adipose tissue (WAT) weights were unaffected by drug treatment (Fig. 2a), but 0602K trended to increase inguinal subcutaneous WAT and significantly increased BAT weight whether provided alone or in combination with Lira (Fig. 2b,c). Increased liver weights in *db/db* mice and not altered by drug treatment (Fig. 2d), yet hepatic TAG was nearly significantly reduced by 0602K+Lira (Fig. 2e). Hepatic glycogen levels were also increased in *db/db* mice and were completely normalized by either 0602K or 0602K+Lira (Fig. 2f). Thus, 0602K increased adiposity and corrected hepatic glycogen levels whether given alone or in combination with Lira.

**Fig. 2.**
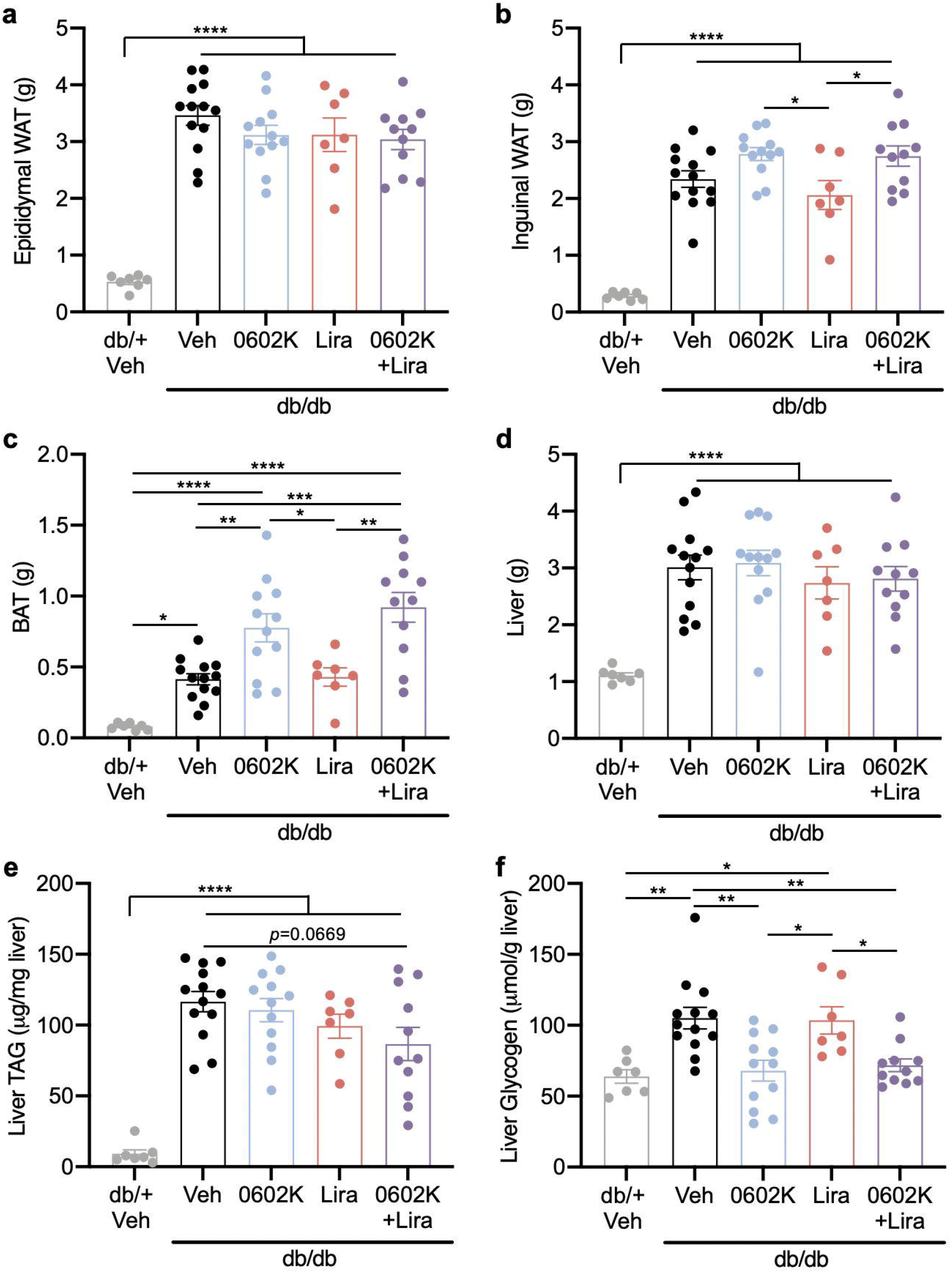
Effects of MSDC-0602K or Liraglutide on tissue weights, liver triglycerides, and glycogen. (**a**-**d**) Epididymal (visceral) white adipose tissue (WAT), inguinal (subcutaneous) WAT, intrascapular brown adipose tissue (BAT) and liver weights. (**e**-**f**) Hepatic triglyceride (TAG) and glycogen concentrations. n= 7 *db/*+ Veh; 13 *db/db* Veh, 12 *db/db* 0602K, 7 *db/db* Lira, and 11 *db/db* 0602K+Lira. All data are expressed measn±SEM within dot plot of individual data points. Individual data points represent a single mouse. Ordinary one-way ANOVA with Tukey’s multiple comparison’s test: \**p*<0.05, \*\**p*<0.01, \*\*\**p*<0.001, and \*\*\*\**p*<0.0001.

### Effects of MSDC-0602K and Liraglutide on insulinemia and plasma lipids

Vehicle-treated *db/db* mice were hyperinsulinemic and 0602K decreased these plasma insulin concentrations (Fig. 3a). In contrast, Lira treatment further increased plasma insulin, yet mice treated with 0602K+Lira displayed reduced insulin similar to 0602K-treated animals (Fig. 3a). To resolve whether the changes in insulinemia were due to insulin secretion versus insulin clearance, we measured plasma C-peptide concentrations, which were strongly elevated in vehicle-treated *db/db* mice. While Lira further increased C-peptide concentrations, 0602K or 0602K+Lira-treated mice displayed reduced C-peptide (Fig. 3b). An increased insulin/C-peptide ratio in 0602K-treated mice suggests that 0602K more strongly reduced insulin secretion versus increased clearance (Fig. 3c). Plasma NEFA, TAG, and cholesterol were all increased in vehicle-treated *db/db* mice, and reduced by 0602K, Lira, or 0602K+Lira treatments (Fig. 3d-f). These results suggest that 0602K reduces hyperinsulinemia by reducing insulin secretion even in combination with Liraglutide, yet all treatments were able to improve plasma lipids.

**Fig. 3.**
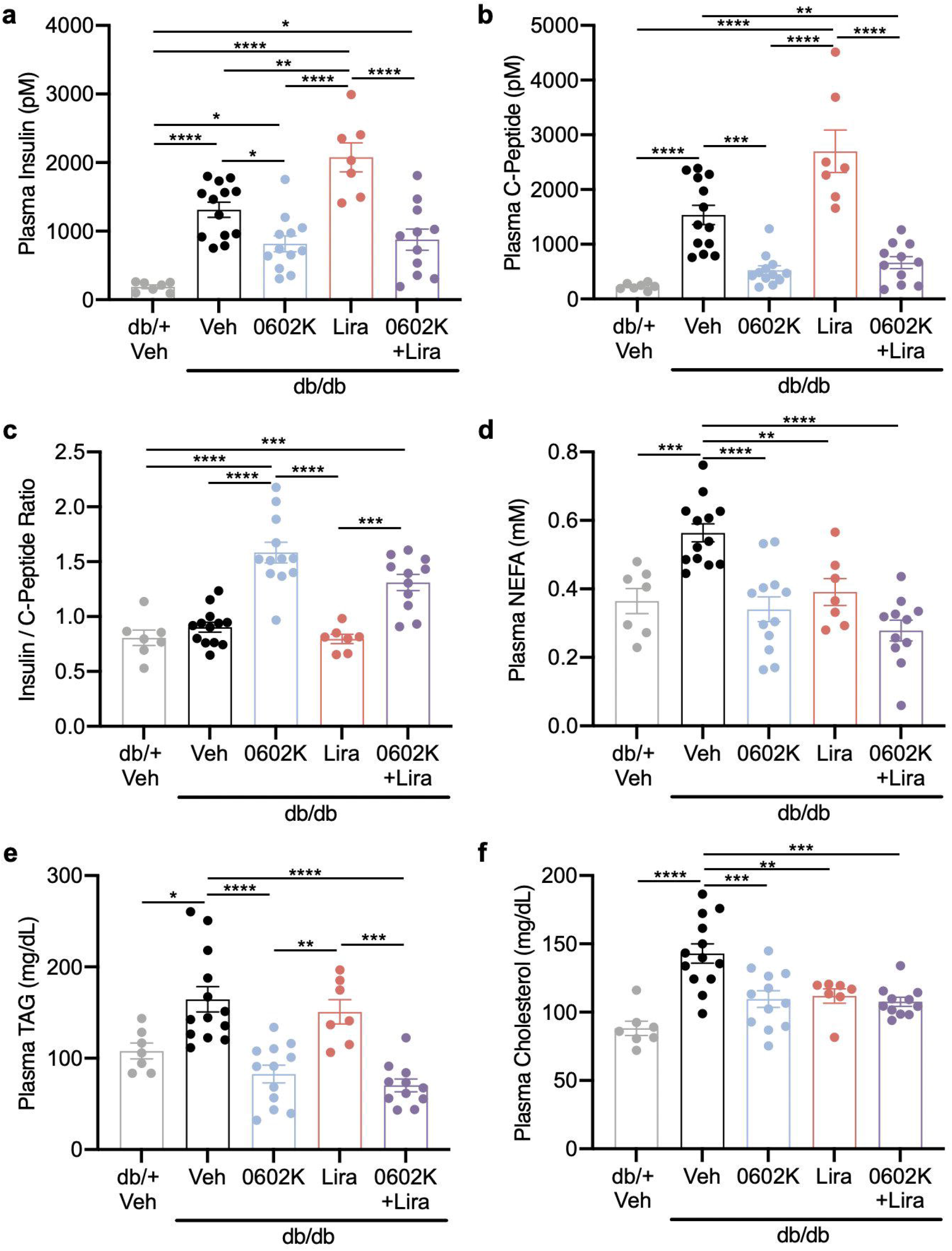
MSDC-0602K reduces insulinemia and plasma lipids. (**a**-**f**) Plasma insulin, C-peptide, insulin/C-peptide ratio, non-esterified fatty acids (NEFA), triglycerides (TAG), and cholesterol. n= 7 *db/*+ Veh; 13 *db/db* Veh, 12 *db/db* 0602K, 7 *db/db* Lira, and 11 *db/db* 0602K+Lira. All data are expressed means±SEM within dot plot of individual data points. Individual data points represent a single mouse. Ordinary one-way ANOVA with Tukey’s multiple comparison’s test: \**p*<0.05, \*\**p*<0.01, \*\*\**p*<0.001, and \*\*\*\**p*<0.0001.

### MSDC-0602K improves islet insulin content by indirectly reducing insulin secretion

Immunofluorescence of pancreas sections showed that the hyperinsulinemia of vehicle-treated *db/db* mice caused a dramatic reduction of islet insulin content (Fig. 4a,b). 0602K-treatment alone or in combination with Lira significantly improved islet insulin content (Fig. 4a,b). We next wanted to test whether 0602K was directly inhibiting insulin secretion. This is particularly intriguing as 0602K inhibits the MPC [9, 10], and we have previously shown that mice with beta cell-specific MPC-deletion display defective GSIS [27]. However, 0602K treatment of isolated islets had no effect on GSIS (Fig. 4c). Lastly, a single gavaged dose of 0602K was able to reduce plasma insulin and C-peptide concentrations in diet-induced obese fl/fl (wildtype) and RIPCre-driven beta cell MPC2 knockout mice (Fig. 4d-f). Thus, insulin secretion was reduced by 0602K even when the molecular target of 0602K was not expressed in the pancreatic beta cells. Altogether, these results suggest that 0602K preserves islet insulin content by indirectly reducing insulin secretion due to improved peripheral insulin sensitivity.

**Fig. 4.**
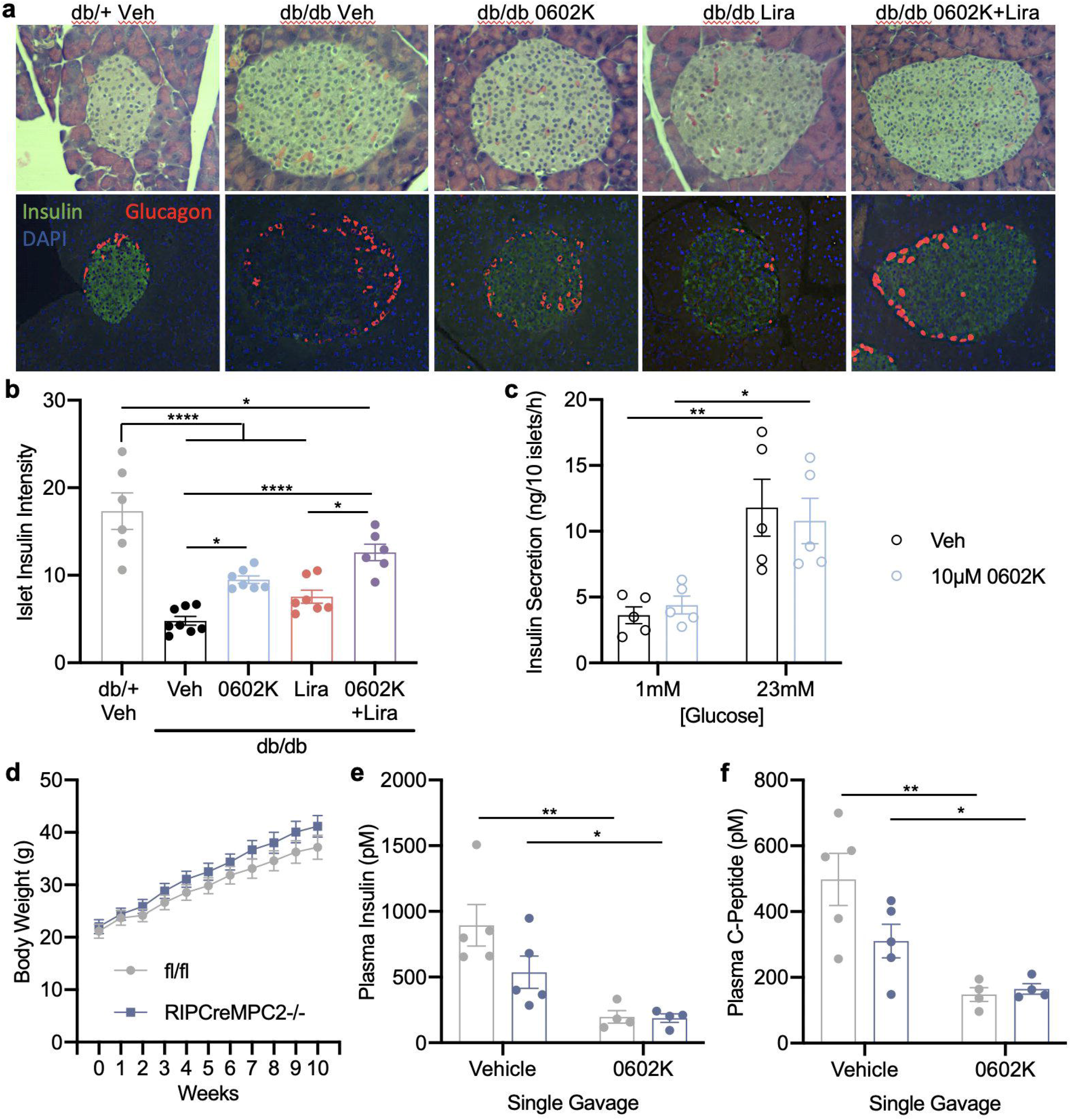
MSDC-0602K indirectly reduces insulin secretion and improves islet insulin content. (**a**) Representative H&E images and insulin/glucagon immunofluorescence of pancreatic islets. (**b**) Islet insulin fluorescence intensity measured by the islet green fluorescence normalized to green fluorescence of surrounding exocrine pancreas tissue. Individual data points represent the average intensity of at least 12 islets per mouse. (**c**) Insulin secretion from isolated wildtype islets at 1 versus 23mM glucose comparing DMSO vehicle with 10μM MSDC-0602K treatment. Individual data points represent the average of an independent experiment, each containing 3-4 individual replicates. (**d**) Average weekly body weights of fl/fl (wildtype) and beta cell specific MPC2 knockout (RIPCreMPC2-/-) mice fed high-fat diet for 10 weeks. After 10 weeks of high-fat diet, mice were gavaged once with vehicle or 30mg/kg MSDC-0602K and euthanized the following morning after ∼16 hours. (**e**,**f**) Plasma insulin and C-peptide measured after a single dose of vehicle or 0602K. All data are expressed as means±SEM or means±SEM within dot plot of individual data points. Individual data points represent a single mouse. Ordinary one-way ANOVA with Tukey’s multiple comparison’s test: \**p*<0.05, \*\**p*<0.01, \*\*\**p*<0.001, and \*\*\*\**p*<0.0001.

### MSDC-0602K and Liraglutide improve liver pathology in a mouse model of NASH

Insulin resistance is a strong driver of NAFLD, and 0602K improves aspects of NASH histology in both mice and humans [13, 16]. To test if NASH was more significantly improved with dual 0602K and Lira therapy, MS-NASH mice were fed a western diet and provided fructose in the drinking water to develop obesity and NASH. Vehicle, 0602K, Lira, or 0602K+Lira treatments were started after 18 weeks on diet, and similar to the *db/db* study, 0602K-treated mice tended to display increased body weight (Fig. 5a,b). Plasma ALT and AST were monitored monthly, and while Lira caused modest reductions in these plasma markers of liver injury, 0602K or 0602K+Lira induced more significant reductions (Fig. 5c-f). Only 0602K+Lira reduced liver weights compared to vehicle-treated mice, however as a percentage of body weight, 0602K- or 0602K+Lira-treated mice displayed reduced liver size (Fig. 6a,b). Liver TAG and glycogen concentrations were unaffected by any treatment in these mice (Fig. 6c,d). Scoring of liver histology agreed with the biochemical measurements that steatosis was not improved by any treatment (Fig. 6e,f). However, hepatic inflammation and ballooning were significantly improved by combination 0602K+Lira treatment (Fig. 6g,h). Combined, while 0602K or Lira alone improved the NAFLD activity score, 0602K+Lira combination more significantly improved NASH liver histology (Fig. 6e,i). Thin bridging fibrosis was identified in all mice, and while the fibrosis histology scores were not improved by any treatment (data not shown), 0602K or 0602K+Lira treatment reduced the expression of several collagen and extracellular matrix-related genes (Fig. 6j). Altogether, these results indicate that combination of 0602K and Lira improves NASH in this mouse model.

**Fig. 5.**
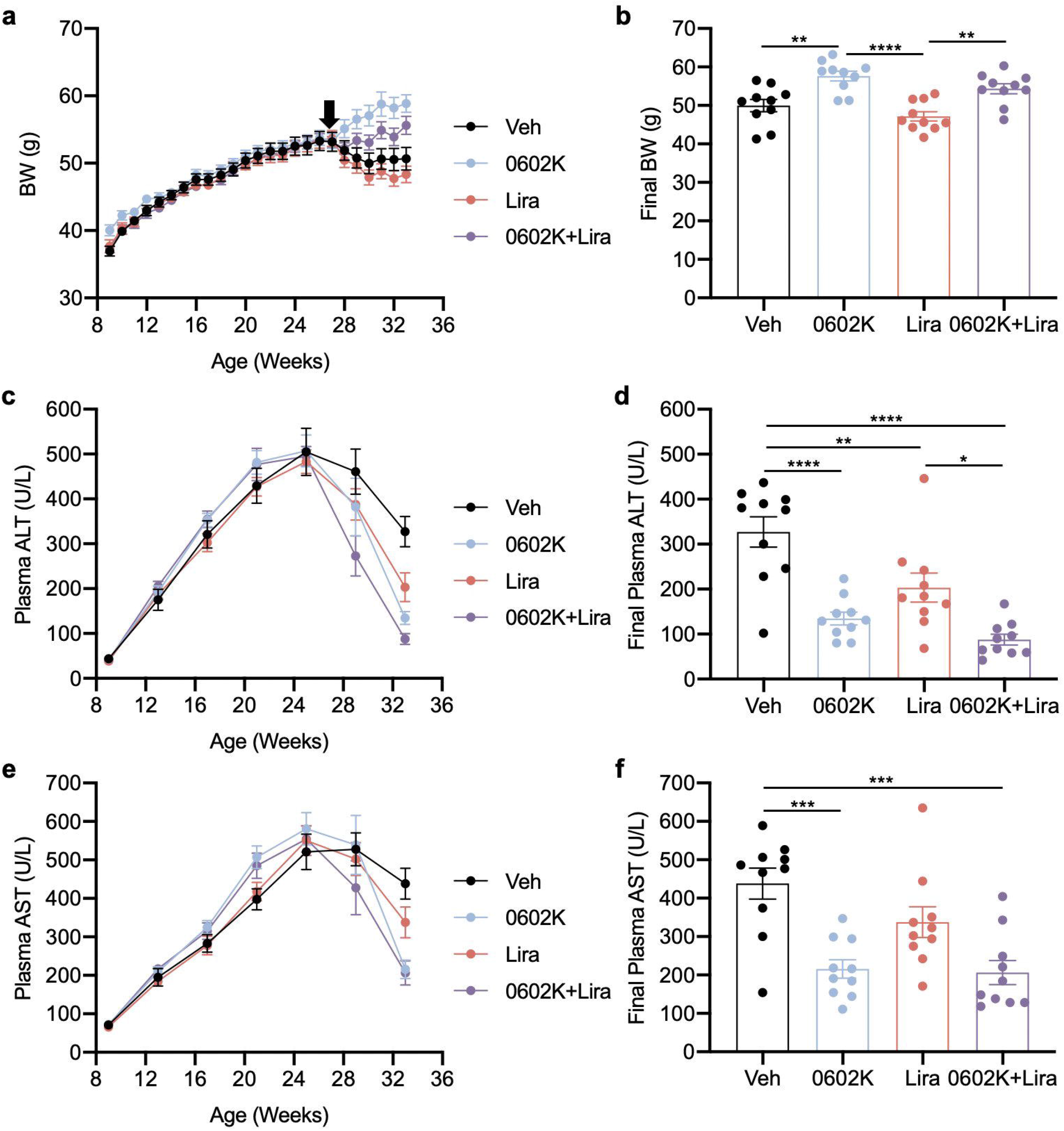
MSDC-0602K and Liraglutide improve plasma measures of liver injury. (**a**,**b**) Average weekly body weight (BW) and final BW of MS-NASH mice treated with either Vehicle, 0602K, Liraglutide, or 0602K+Lira. (**c**-**f**) Average monthly plasma alanine aminotransferase (ALT) and aspartate aminotransferade (AST) and final ALT and AST. n=10 for all groups. All data curves are expressed as means±SEM, and final data expressed as means±SEM within dot plot of individual data points. Individual data points represent a single mouse. Ordinary one-way ANOVA with Tukey’s multiple comparison’s test: \**p*<0.05, \*\**p*<0.01, \*\*\**p*<0.001, and \*\*\*\**p*<0.0001.

**Fig. 6.**
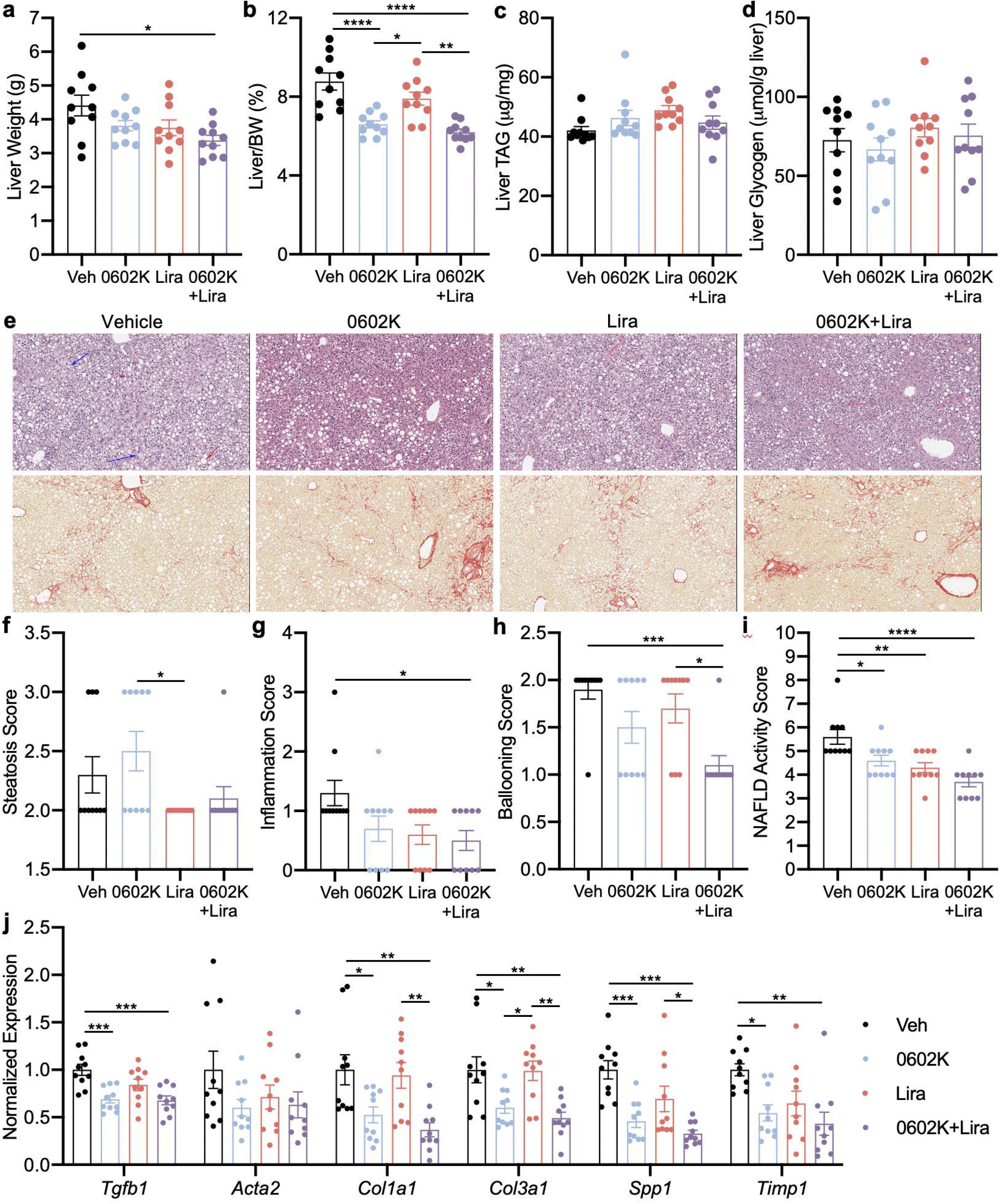
MSDC-0602K+Liraglutide treatment improves NASH liver pathology. (**a**,**b**) Liver weight and liver weight normalized to body weight (BW). (**c**,**d**) Liver triglyceride (TAG) and glycogen concentrations. (**e**) Representative liver H&E and Sirius Red histology images. (**f**-**i**) Liver histology scoring for NAFLD (steatosis, inflammation, hepatocyte ballooning, and combined NAFLD activity score) determined by a histopathologist blinded to treatment groups. (**j**) Hepatic gene expression for genes related to hepatic stellate cell activation, collagen, and extracellular matrix organization. n=10 for all groups. All data expressed as means±SEM within dot plot of individual data points. Individual data points represent a single mouse. Ordinary one-way ANOVA with Tukey’s multiple comparison’s test: \**p*<0.05, \*\**p*<0.01, \*\*\**p*<0.001, and \*\*\*\**p*<0.0001.

### MSDC-0602K decreases insulinemia and improves pancreas insulin content in MS-NASH mice

0602K, Lira, or 0602K+Lira treatments did not reduce blood glucose in these MS-NASH mice (Fig. 7a). However, 0602K or 0602K+Lira treatments strongly reduced plasma insulin (Fig. 7b). Plasma NEFA were also significantly reduced by 0602K or 0602K+Lira treatment (Fig. 7c), suggesting improved insulin action and decreased adipose lipolysis. Lastly, insulin immunohistochemistry identified a greater number of insulin+ cells in 0602K or 0602K+Lira treated pancreata (Fig. 7d,e). Thus, similar to the *db/db* study, these results suggest that the improved insulin sensitivity with 0602K increases islet insulin content.

**Fig. 7.**
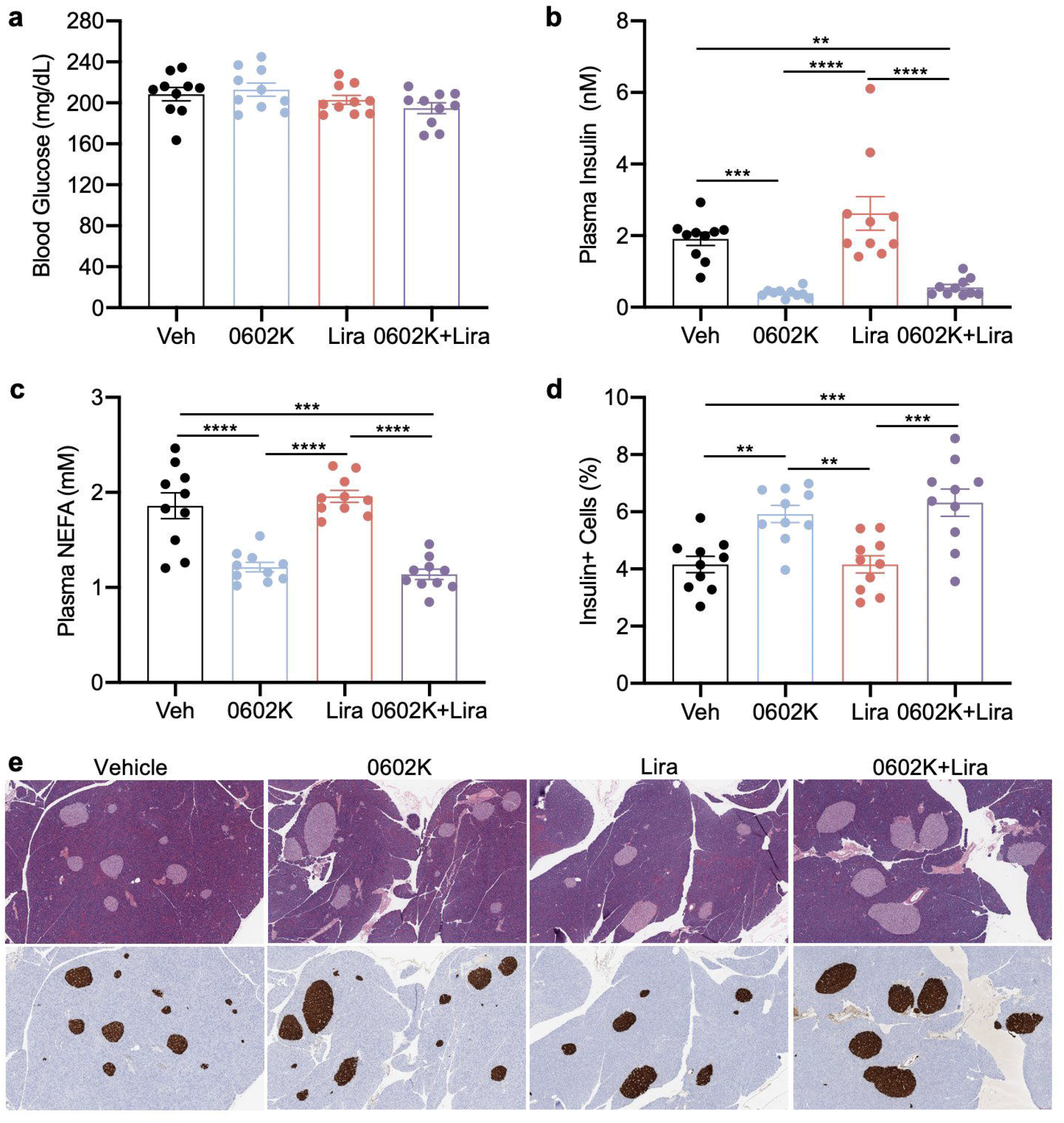
MSDC-0602K improves insulinemia and islet pancreatic insulin content in MS-NASH mice. (**a**-**c**) Blood glucose concentrations, plasma insulin concentrations, and plasma non-esterified fatty acids (NEFA). (**d**,**e**) Percentage of insulin positive cells in pancreas sections and representative pancreas H&E histology and insulin immunohistochemistry images. n=10 for all groups. All data expressed as means±SEM within dot plot of individual data points. Individual data points represent a single mouse. Ordinary one-way ANOVA with Tukey’s multiple comparison’s test: \**p*<0.05, \*\**p*<0.01, \*\*\**p*<0.001, and \*\*\*\**p*<0.0001.

## Discussion

The overall goal of these present studies was to assess whether combination of a novel TZD insulin sensitizer and a GLP-1RA would improve diabetes and NASH better than either individual therapy. Studies in *db/db* mice suggested that combining MSDC-0602K and Liraglutide did indeed provide more significant improvement in glucose tolerance. Glycemia and insulinemia were greatly improved with 0602K and not further improved by 0602K+Lira treatment. 0602K also improved pancreatic insulin content, and 0602K+Lira further increased islet insulin content. In MS-NASH mice, while plasma biomarkers and some aspects of NASH histology were improved by each treatment, more significant improvements were achieved with combination 0602K+Lira. A secondary goal of these studies was to test if the weight loss associated with Liraglutide could prevent the weight gain associated with TZDs. In the *db/db* study, 0602K+Lira treated mice displayed similar slight weight gain as 0602K-treated mice, however there was some attenuation of weight gain from combined 0602K+Lira treatment in the MS-NASH study. A study in diabetic rats using a rather low dose of Pioglitazone and double the dose of Lira as used in the current study also observed the greatest improvement in glucose tolerance with Pioglitazone+Lira, and while Lira-treated rats lost weight, Pioglitazone+Lira did not attenuate the Pioglitazone-induced weight gain [31]. However, a study of Pioglitazone+Lira in *db/db* mice did observe attenuation of the Pioglitazone-induced weight gain with combined Lira treatment [32].

The strongest effects observed in our *db/db* study were the complete correction of glycemia and reduction in insulinemia with 0602K. This decrease in insulinemia with TZDs has been well described in both humans and animal models of insulin resistance and diabetes [33-35]. The presiding dogma is that improved insulin sensitivity indirectly reduces the need for insulin hyper-secretion, and thus TZDs can improve beta cell insulin content and function [3, 35-37]. However, it has recently been recognized that traditional and PPARγ-sparing TZDs can bind and inhibit the MPC [8-10], and pyruvate import into the mitochondria via the MPC plays an important role in beta cell glucose sensing and GSIS [27, 38]. This raises the possibility that MPC-inhibition with TZDs may directly inhibit beta cell insulin secretion, and indeed there are reports of acute TZD treatment decreasing GSIS [39, 40]. In this current study however, MSDC-0602K did not reduce GSIS in isolated islets (Fig. 4c), and 0602K was still able to reduce plasma insulin and C-peptide concentrations in mice lacking MPC expression in beta cells (Fig. 4e,f). A similar TZD molecule, MSDC-0160, was also found to have no effect on GSIS but improved insulin content and beta cell phenotype in isolated human islets [41]. Interestingly, MPC-inhibition with the tool-compound UK-5099 appears to alleviate the metabolic stress of islets cultured in high glucose [42]. Lastly, the preserved islet insulin content with TZDs can lead to enhanced insulin secretion during glucose challenge [35, 36]. Altogether, these findings suggest that TZDs are not directly inhibiting GSIS, rather improved peripheral insulin sensitivity indirectly reduces the need for excessive insulin secretion.

While we observed similar decreases in insulinemia with 0602K or 0602K+Lira in the MS-NASH model of nonalcoholic steatohepatitis, the main goal of this experiment of course was to analyze fatty liver pathology. All treatments improved plasma ALT and AST, as well as certain aspects of liver histology, with the combination of 0602K+Lira typically resulting in more significant improvements (Fig. 5c-f, Fig. 6e-i). Similar to our previous study in which we fed C57BL/6J mice a NASH-inducing diet [16], 0602K treatment did not improve hepatic triglycerides and steatosis (Fig. 6c,e,f). Conversely, steatosis was improved by 0602K in humans with NASH [13]. Our previous study identified reductions in histologic fibrosis with 0602K [16], which were not observed in the current model, even with 0602K+Lira combination. Yet both the current and previous studies identified reduced expression of genes related to hepatic stellate cell activation and fibrosis, and these were not further reduced by 0602K+Lira (Fig. 6j).

There were a limited number of analyses that displayed additive improvement with combined 0602K+Lira treatment. Namely, glucose tolerance in the *db/db* mice and liver weights and combined NAFLD activity scores in the MS-NASH mice were the main endpoints for which 0602K+Lira outperformed 0602K-only treatment. This could be due to a rather low dose of Liraglutide used in these studies (0.2mg/kg every-other-day), compared to other rodent studies that use similar doses with daily or even twice daily injections [32, 43, 44]. However, we are confident that we provided an effective dose of Liraglutide as Lira-treated animals displayed weight loss, increased plasma insulin concentrations, and improved glycemia. Most surprisingly, even though Lira increased insulin secretion, combination with 0602K resulted in similar suppression of insulinemia compared to 0602K-treated mice with improved effects on glucose tolerance.

In conclusion, while MSDC-0602K improved many aspects of insulin resistance and NASH by itself, glucose tolerance and several aspects of NASH were more significantly improved by combining 0602K and Liraglutide. In the MS-NASH study, Lira attenuated the weight gain associated with 0602K, but this was not the case in *db/db* mice. This study also clarifies that TZDs likely reduce insulinemia not by directly inhibiting the MPC in beta cells but indirectly by improving peripheral insulin sensitivity. It remains unclear why MPC-inhibition with TZDs does not reduce GSIS unlike genetic MPC deletion [16]. Nonetheless, these studies suggest that combining insulin sensitizer and GLP-1RA therapies may better improve diabetes and fatty liver disease and potentially attenuate TZD-induced weight gain.

### Data availability

All data generated during these studies are included in the text, figures, and tables of this article and electronic supplementary material. Source data or materials will be supplied by the corresponding author with reasonable request.

## Funding

K.S.M. is supported by NIH R00 HL136658. The *db/db* study and analyses received no specific grant funding from any public or commercial agency. The MS-NASH mouse study was performed at Crown Biosciences Inc. and was funded by Cirius Therapeutics.

## Authors’ relationships and activities

J.R.C. is an employee, chief scientific officer, and stockholder of Cirius Therapeutics which is developing MSDC-0602K for NASH. All other authors declare no relationships or conflicts of interest.

## Contribution statement

D.R.K., K.D.P., and M.S. performed the experiments, analyzed the data, assisted in manuscript writing and editing. L.H. analyzed the liver histology slides. J.R.C. conceived and designed the research, and wrote and edited the manuscript. K.S.M. conceived and designed the research, performed experiments, analyzed the data, and wrote and edited the manuscript. K.S.M. is the guarantor of this work and accepts full responsibility for the conduct of the performance and analyses of these studies, maintains full access to the data, and controlled the decision to publish.

