## Supplemental Table 1 for "MSDC-0602K, A Novel Insulin-Sensitizer Improves Insulinemia and Fatty Liver Disease Alone and in Addition to Liraglutide in Mice"

**Supplementary Table 1. Primer oligonucleotide sequences used for qRT-PCR of mouse liver.**

| Gene | Forward 5'-3' | Reverse 5'-3' | Amplicon Length |
|---|---|---|---|
| *Acta2* | gtc cca gac atc agg gag taa | tcg gat act tca gcg tca gga | 102 |
| *Col1a1* | gct cct ctt agg ggc cac t | cca cgt ctc acc att ggg g | 103 |
| *Col1a3* | ctg taa cat gga aac tgg gga aa | cca tag ctg aac tga aaa cca cc | 144 |
| *Rplp0* | gca gac aac gtg ggc tcc aag cag at | ggt cct cct tgg tga aca cga agc cc | 190 |
| *Spp1* | atc tca cca ttc gga tga gtc t | tgt agg gac gat tgg agt gaa a | 79 |
| *Tgfb1* | ctc ccg tgg ctt cta gtg c | gcc tta gtt tgg aca gga tct g | 133 |
| *Timp1* | cca gag ccg tca ctt tgc tt | agg aaa agt aga cag tgt tca ggc tt | 126 |
